# A facile platform to engineer *E. coli* tyrosyl-tRNA synthetase adds new chemistries to the eukaryotic genetic code, including a phosphotyrosine mimic

**DOI:** 10.1101/2021.11.28.470256

**Authors:** Katherine T. Grasso, Soumya Jyoti Singha Roy, Megan Jin Rae Yeo, Chintan Soni, Arianna O. Osgood, Christen M. Hillenbrand, Abhishek Chatterjee

## Abstract

The *E. coli* tyrosyl-tRNA synthetase (EcTyrRS)/tRNA^EcTyr^ pair offers an attractive platform to genetically encode new noncanonical amino acids (ncAA) in eukaryotes. However, challenges associated with a eukaryotic selection system, which is needed for its engineering, has impeded its success in the past. Recently, we showed that EcTyrRS can be engineered using a facile *E. coli* based selection system, in a strain where the endogenous tyrosyl pair has been substituted with an archaeal counterpart. However, a significant cross-reactivity between the UAG-suppressing tRNA_CUA_^EcTyr^ and the bacterial glutaminyl-tRNA synthetase limited the scope of this strategy, preventing the selection of moderately active EcTyrRS mutants. Here we report an engineered tRNA_CUA_^EcTyr^ that overcomes this cross-reactivity. Optimized selection systems using this tRNA enabled efficient enrichment of both strongly and weakly active ncAA-selective EcTyrRS mutants. We also developed a wide-dynamic range (WiDR) antibiotic selection to further enhance the activities of the weaker first-generation EcTyrRS mutants. We demonstrated the utility of our platform by developing several new EcTyrRS mutants that efficiently incorporate useful ncAAs in mammalian cells, including photo-affinity probes, bioconjugation handles, and a non-hydrolyzable mimic of phosphotyrosine.

## Introduction

Site-specific incorporation of noncanonical amino acids (ncAAs) into proteins in living cells is a popular technology with numerous enabling applications.^1–5^ The ncAA is typically encoded using a repurposed nonsense codon, and an engineered nonsense-suppressing aminoacyl-tRNA synthetase (aaRS)/tRNA pairs is used to deliver it during translation. To maintain the fidelity of translation, the ncAA-selective aaRS/tRNA pair must not cross-react with its host counterparts. Typically, such an ‘orthogonal’ pair is imported into the host cell from a different domain of life. For example, bacteria-derived pairs are typically suitable for ncAA incorporation in eukaryotes, while eukaryotic and archaea derived pairs are used in bacteria.^1, 3–5^ Although multiple bacteria-derived pairs have been developed for such application in eukaryotes,^6–10^ the ncAA toolbox therein has been disproportionately reliant on the unique archaea-derived pyrrolysyl pair, which is orthogonal across prokaryotes and eukaryotes.^1–4, 11^ The success of this pair stems partly from the innate plasticity of the pyrrolysyl-tRNA synthetase (PylRS), but another major advantage is the ability to engineer the PylRS using a facile *E. coli* based selection system.^4^ In contrast, other eukaryote-compatible aaRS/tRNA pairs that are derived from bacteria, must be engineered using a cumbersome yeast-based selection system.^6, 8^ Despite its remarkable success, the pyrrolysyl pair provides access to only certain structural classes of ncAAs. Access to additional readily evolvable pairs is critical to expand the structural diversity of the genetically encoded ncAAs in eukaryotic cells.

The *E. coli* derived tyrosyl-tRNA synthetase (EcTyrRS)/tRNA^EcTyr^ pair was first established for ncAA incorporation in eukaryotes nearly two decades ago.^6–7^ Engineering this pair can provide access to ncAAs that are currently unavailable for incorporation in eukaryotes, such as those mimicking polar post-translational modifications of tyrosine.^10, 12^ Indeed, the archaea-derived TyrRS/tRNA pair, which has a similar active site architecture to its bacterial counterpart, has been engineered to encode over a hundred different ncAAs in prokaryotes,^2–3^ but cannot be used in eukaryotes due to cross-reactivity. Its success highlights the untapped potential of the (EcTyrRS)/tRNA^EcTyr^ pair for building a comparable toolbox for application in eukaryotic cells. However, only a small set of ncAAs have been genetically encoded using this platform to date, which can be partially attributed to the challenges associated with the yeast-based selection system needed to engineer it.

To overcome this limitation, we recently developed novel ‘altered translational machinery tyrosyl’ (ATMY) *E. coli* strains, in which the endogenous EcTyrRS/tRNA pair was functionally substituted with an archaeal counterpart (**Figure S1**).^10, 13^ We further demonstrated that the ‘liberated’ EcTyrRS/tRNA^EcTyr^ pair can be reintroduced into ATMY strain as a nonsense suppressor, and its substrate specificity can be engineered using the facile *E. coli* based selection system. Although we have demonstrated that this platform can be used to genetically encode previously inaccessible ncAAs in eukaryotes, such as p-boronophenylalanine and O-sulfotyrosine, ^10, 12^ its scope was significantly limited by significant cross-charging of tRNA_CUA_^EcTyr^ by the endogenous glutaminyl-tRNA synthetase (GlnRS).^10^ Here we show that this cross-reactivity compromises the ability to identify moderately active EcTyrRS mutants from naïve mutant libraries using the first-generation selection system. To overcome this limitation, we used directed evolution to develop an engineered tRNA_CUA_^EcTyr^ that lacks this crossreactivity. An optimized selection system using this tRNA_CUA_^EcTyr^ enabled facile enrichment of ncAA-selective EcTyrRS mutants from naïve libraries, including those with weaker activities. We further developed a selection strategy to improve the performance of first-generation EcTyrRS mutants with weaker activities to create robust ncAA incorporation platforms in mammalian cells. This optimized selection system was used to developed several different EcTyrRS mutants: 1) one that charges p-benzoylphenylalanine (pBPA **1**, **Figure 1**), a popular photoaffinity probe,^6, 14^ with high efficiency in mammalian cells, 2) a highly polyspecific mutant that enables incorporation of a wide range of structurally similar ncAAs with useful handles (e.g., bioconjugation, electrophilic groups that enable proximity-dependent cross-linking, etc.), and 3) a unique mutant that enables the incorporation of p-carboxymethylphenylalanine (pCMF **12**), a non-hydrolyzable mimic of phosphotyrosine.

**Figure 1:**
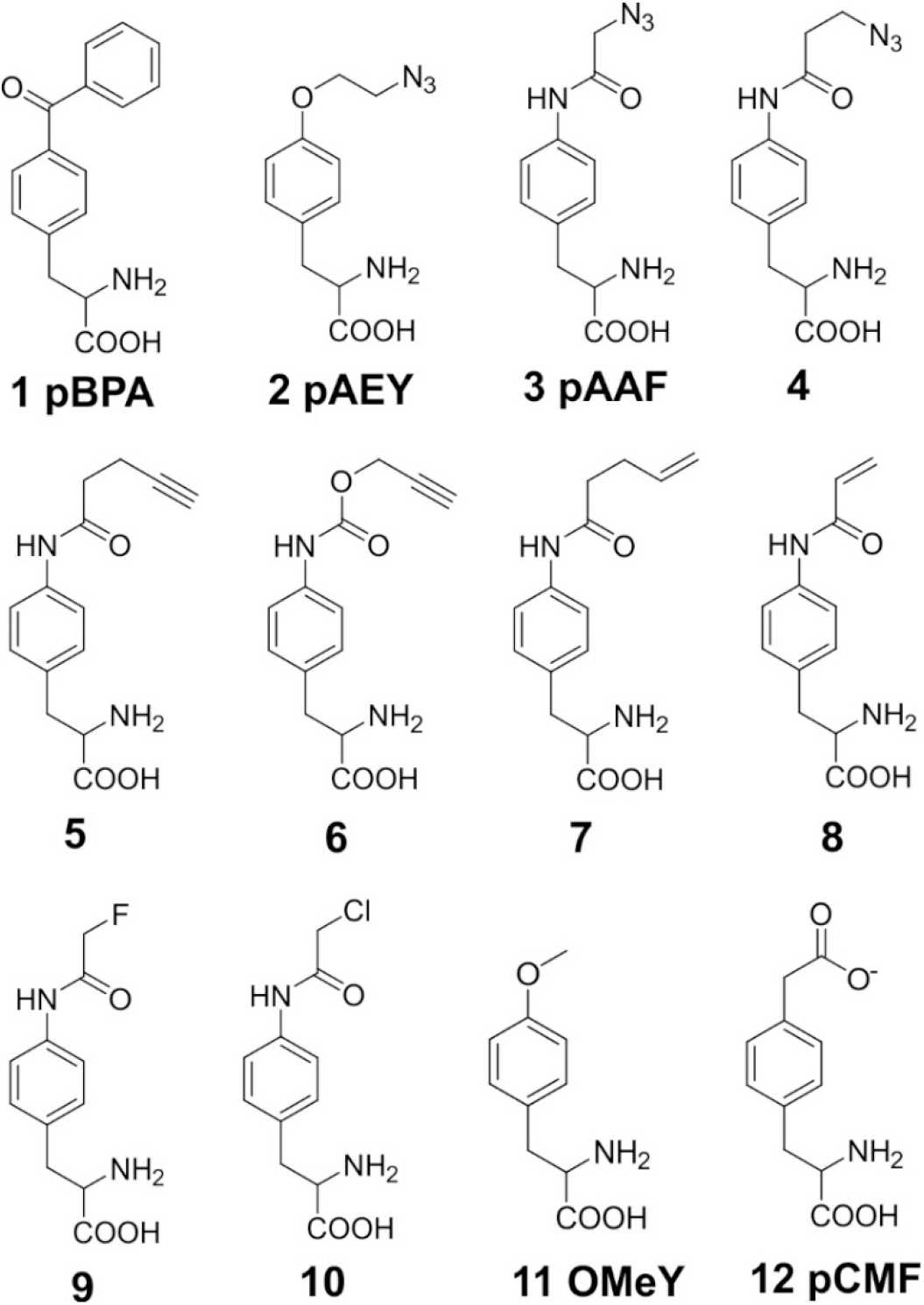
Structures of ncAAs used in this study.

## Results and Discussion

### Development of a mutant tRNA_CUA_^EcTyr^ orthogonal in ATMY *E. coli*

In our ATMY *E. coli* strains, the endogenous tyrosyl pair is replaced with an archaeal counterpart; consequently, these strains are suitable hosts for engineering the EcTyrRS/tRNA^EcTyr^ pair (**Figure S1**).^10^ The first step in developing this directed evolution platform involves establishing the EcTyrRS/tRNA^EcTyr^ pair in the ATMY strain as an active but orthogonal nonsense suppressor. However, we found previously that *E. coli* GlnRS weakly recognizes the TAG-suppressing tRNA_CUA_^EcTyr^ mutant.^10^ This cross-reactivity is driven largely by the well-established affinity *E. coli* GlnRS shows for the engineered CUA anticodon.^9–10, 15–16^ Background activity from this unexpected cross-charging interferes with the selection of ncAA-specific EcTyrRS variants. Previously, we reduced the impact of this cross-reactivity by optimizing the relative expression levels of tRNA_CUA_^EcTyr^ and tRNA^Gln^: 1) A strain (ATMY3) was developed with a single genomic copy of the tRNA_CUA_^EcTyr^ to minimize its expression level, while 2) tRNA^Gln^ was overexpressed from a plasmid to outcompete tRNA_CUA_^EcTyr^ from being charged by GlnRS.^10, 16^ The resulting reduction in cross-reactivity allowed us to establish a directed evolution platform to engineer EcTyrRS in ATMY3, and develop mutants that charge previously inaccessible ncAAs in eukaryotic cells.^10, 12^ However, we soon realized that even though our first-generation selection system can discriminate highly active EcTyrRS mutants from the background cross-activity to facilitate their enrichment, it is unable to do the same for less efficient EcTyrRS mutants. For example, when we tested a previously established EcTyrRS mutant that charge pBPA **1** (pBPARS)^6^ using our selection system, which couples the activity of the EcTyrRS mutant to the expression of a chloramphenicol-acetyl transferase (CAT-98-TAG) reporter, the cells survived up to 10 μg/mL chloramphenicol in the absence of pBPA, and only 15 μg/mL in its presence (**Figure S2**). Such a small difference between the background and the target activity is inadequate to serve as the basis of an effective selection system. It is important to note that the ability to identify moderately active first-generation aaRS mutants is critically important for genetically encoding ncAAs with challenging structural features, for which it can be difficult to directly obtain a highly efficient aaRS mutant from a naïve library. In such cases, the moderately active first-generation mutants can serve as ‘stepping stones’ to develop more efficient variants through further evolution.^17–20^ To enable such engineering efforts on EcTyrRS, we sought to develop a truly orthogonal tRNA_CUA_^EcTyr^ mutant that will serve as the foundation for a more versatile selection system.

The acceptor stem of tRNAs has been often targeted to improve their orthogonality/activity with much success.^21–24^ We fully randomized 5 base pairs in the acceptor stem of tRNA_CUA_^EcTyr^ to generate a library of roughly 10^6^ mutants (**Figure 2A**). The first base pair in the acceptor stem was not randomized, given it is an important identity element. This library was subjected to a round of positive selection, in the presence of a EcTyrRS mutant that we previously developed to charge O-methyltyrosine.^10^ In this step, active tRNA_CUA_^EcTyr^ mutants are enriched based on their ability to express a TAG-inactivated CAT reporter. Next, these mutants are subjected to a negative selection in the absence of a cognate EcTyrRS, where cross-reactive tRNA_CUA_^EcTyr^ mutants are eliminated through the expression of a TAG-inactivated toxic gene (barnase). After two rounds of positive and negative selection, the surviving tRNA_CUA_^EcTyr^ clones were screened for activity and orthogonality. One particular clone (tRNA_CUA_^EcTyr^-h1; **Figure 2B**) was highly promising: When tested with a pBPARS, this tRNA supported growth at up to 30 μg/mL of chloramphenicol in the presence of pBPA, but not even at 5 μg/mL (lowest concentrations tested) in its absence (**Figure 2C**). In contrast, the original tRNA_CUA_^EcTyr^ supported growth at up to 60 μg/mL of chloramphenicol both in the presence and absence of pBPA. Note that this assay did not include the overexpression of tRNA^Gln^, which is necessary to suppress the crossreactivity of the original tRNA_CUA_^EcTyr^, further underscoring the superior orthogonality of tRNA_CUA_^EcTyr^-h1. Interestingly, in addition to a more A-U rich acceptor stem, tRNA_CUA_^EcTyr^-h1 encodes an additional fortuitous G24A mutation in the D-stem (**Figure 2B**).

**Figure 2:**
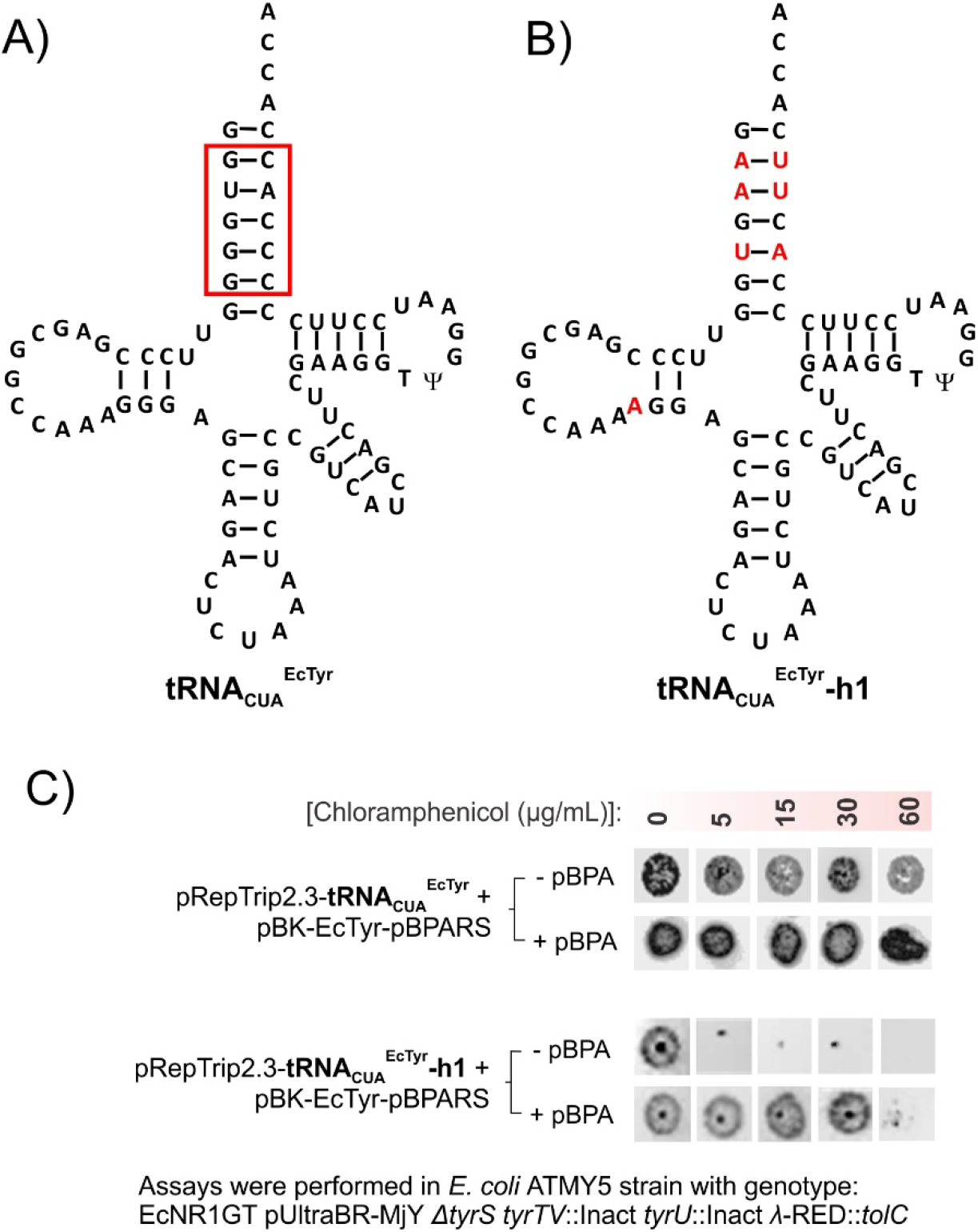
Development of an orthogonal tRNA_CUA_^EcTyr^ variant through directed evolution. A) The sequence of the original tRNA_CUA_^EcTyr^, where the highlighted segment in the acceptor stem was randomized to all possible combinations. B) Selection of this library of mutants led to the identification of tRNA_CUA_^EcTyr^-h1, where the mutations relative to its precursor are shown in red, that shows dramatically attenuated cross-reactivity in ATMY. C) These assays are performed in ATMY5 strain, that does not encode any tRNA_CUA_^EcTyr^, which is instead expressed from the pRepTrip2.3 plasmid, also harboring the CAT-TAG reporter. pBPARS is used as the cognate EcTyrRS and activity is measured in the presence and absence of pBpA. These assays are performed without overexpressing tRNA^Gln^, necessary to reduce the cross-reactivity of original tRNA_CUA_^EcTyr^. Consequently, the tRNA_CUA_^EcTyr^ shows significantly higher background activity (survival at up to 60 μg/mL of chloramphenicol) in the absence of pBPA, which cannot be differentiated from the activity of pBPARS in the presence of pBPA (top panel). In contrast, tRNA_CUA_^EcTyr^-h1 shows no activity in the absence of pBpA, and survival up to 30 μg/mL in the presence of pBPA, showcasing its high orthogonality, and the ability to adequately discern the weak activity of pBPARS from background.

Next, we sought to demonstrate that a selection system employing the orthogonal tRNA_CUA_^EcTyr^-h1 can enable the enrichment of pBPA-selective mutants from a naïve EcTyrRS library, which we were unable to accomplish using the first-generation selection system. A library of ~10^7^ EcTyrRS mutants were constructed by randomizing six residues in the EcTyrRS active site (**Figure 3A**). Additionally, the previously described beneficial mutation D265R in the anticodon binding domain,^10, 25^ which presumably enhances interaction with the non-native CUA anticodon of the cognate tRNA, was included in this library as well. Subjecting this library to an established double-sieve selection system, led to the identification of three clones, including the previously reported pBPARS-1,^6^ that selectively charge pBPA with comparable efficiency (**Figure 3A-B**). While these results confirmed that the new selection system is capable of enriching weakly active EcTyrRS mutants, the poor efficiency of the pBPARS mutants underscores the need to develop a strategy to further improve the activity of such first-generation mutants. A more efficient mammalian incorporation system for pBPA in particular is expected to be useful, given its popularity as an efficient photo-affinity probe.^14, 26–31^

**Figure 3:**
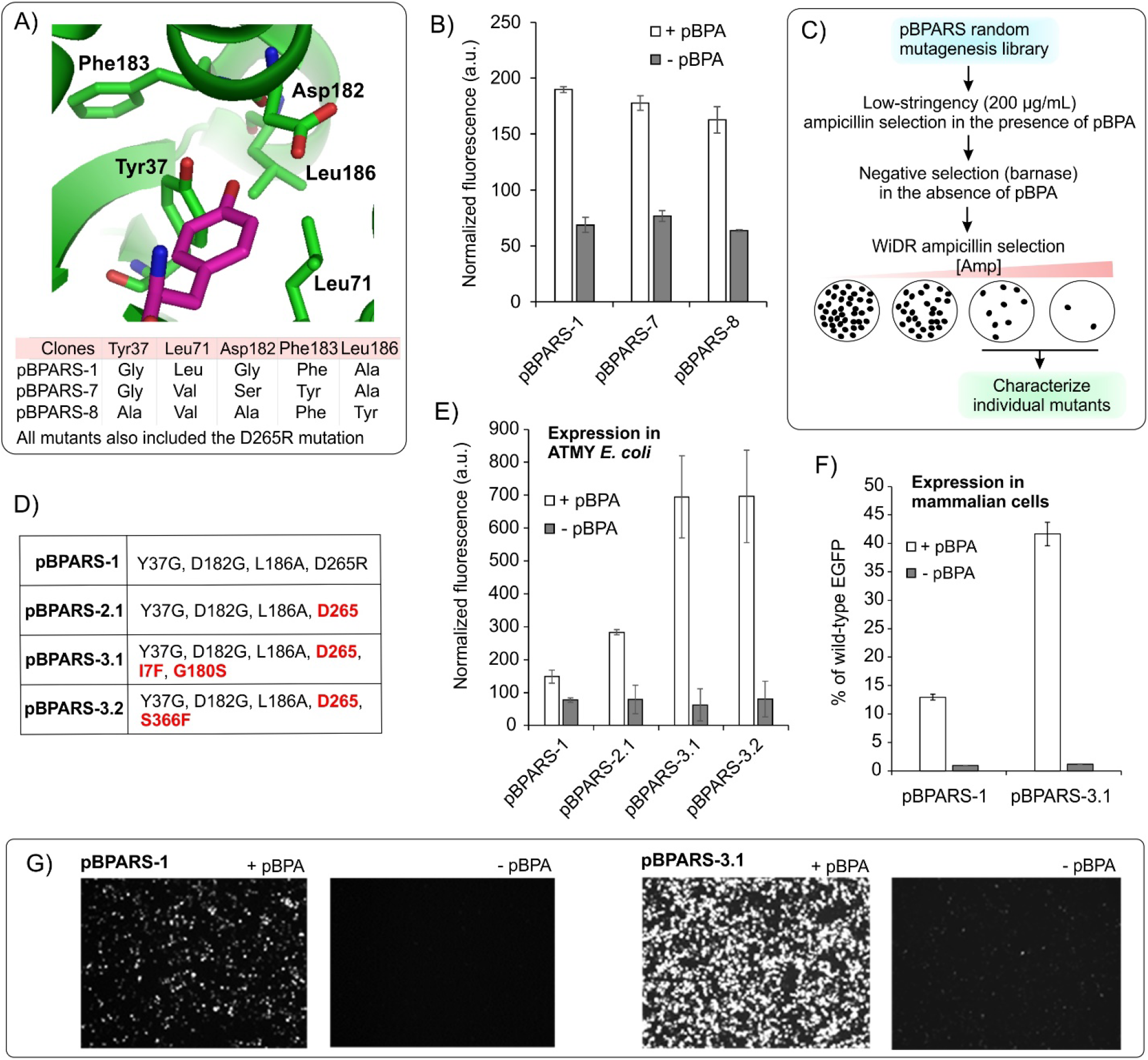
Directed evolution of highly active pBPA-selective EcTyrRS mutants. A) The active site of EcTyrRS with the bound substrate shown in magenta, and the key active site residues that are subjected to randomization highlighted. Mutations in the isolated pBPARS clones are also shown below. B) Activity of the pBPARS mutants in the ATMY E. coli strain, measured through the expression of the sfGFP-151-TAG reporter. C) The scheme for WiDR antibiotic selection to identify pBPARS mutants with higher activity from a random mutagenesis library. D) Mutations associated with enhanced pBPARS mutants (shown in red). E) Activity of the new pBPARS mutants in the ATMY *E. coli* strain, measured through the expression of the sfGFP-151-TAG reporter. F) Activity of the pBPARS mutants in the HEK293T cells, measured through the expression of the EGFP-39-TAG reporter. G) Representative fluorescence images of HEK293T cells expressing the EGFP-39-TAG reporter from panel F.

To evolve next-generation pBPARS mutants with enhanced activity, we needed a selection system that can discriminate between mutants exhibiting moderate and high activity. Although fluorescent protein reporters have been used in the past for this purpose,^19–20^ such selections typically have a lower throughput. In contrast, survival selections using antibiotic reporters are straightforward and provide significantly higher throughput, but have not been used for evolving highly active aaRS mutants from weaker precursors. To explore if an antibiotic-based survival selection is suitable for this purpose, we used two known EcTyrRS mutants that charge O-methyltyrosine (OMeYRS) either weakly or efficiently (**Figure S3A**).^10^ We found that the CAT-based selection system, which has been traditionally used for aaRS evolution, is not able to adequately discriminate between these two EcTyrRS mutants (**Figure S3B**). A screen for additional antibiotic resistance reporters offering a wider dynamic range (WiDR) identified a TAG-inactivated β-lactamase reporter that allows the survival of the strong OMeYRS at up to 1200 μg/mL of ampicillin, but only 400 μg/mL for the weak OMeYRS (**Figure S3B**). This WiDR selection window provided an opportunity to rapidly screen libraries of pBPARS mutants to identify more active variants. Subjecting a library of pBPARS-1 mutants, constructed through error-prone PCR, to this WiDR antibiotic selection (**Figure 3C**) led to the identification of several clones with roughly two-fold higher activity (**Figure 3D-E**), wherein the 265 position reverted back to Asp (pBPARS-2.1). Another round of WiDR evolution using pBPARS-2.1 as the starting point resulted in two new clones with a further ~2-2.5-fold increase in activity (pBPARS-3.1 and 3.2; **Figure 3D-E**). Further evaluation in mammalian cells (**Figure 3F**) revealed that the pBPARS-3.1 mutant facilitates significantly improved pBPA incorporation (>40% of wild-type reporter expression) relative to the original pBPARS-1 (<15% of wild-type reporter expression). These results show that our optimized selection scheme now enables the identification of weakly active EcTyrRS mutants, and that the activity such mutants can be subsequently improved upon using the WiDR antibiotic selection. In addition, the development of pBPARS-3.1, which enables remarkably more efficient incorporation of pBPA in mammalian cells relative to the existing incorporation system, would significantly enhance the utility of this useful photoaffinity probe.

We also explored the mechanism underlying the improved activity of pBPARS-3.1. Reverting back either I7F and G180S that appear in pBPARS-3.1 (**Figure S4A**) were found to be detrimental, suggesting that both mutations contribute to the enhanced activity (**Figure S4B**). We recently reported that mutants of EcTyrRS often exhibit low thermostability. In particular, pBPARS-1 was found to be largely insoluble under ambient conditions, which likely explains its poor activity. Interestingly, we found that a large fraction of pBPARS-3.1 is found in the soluble fraction of E. coli cell-free extract, while nearly all of pBPARS-1 is found in the insoluble fraction (**Figure S4C**). A thermostability assay in *E. coli* cell-free extract further showed that pBPARS-3.1 exhibits significantly higher thermostability (soluble at up to 50 °C) relative to pBPARS-1 (insoluble even at ambient temperature) (**Figure S4D**). These observations suggest that mutations acquired during directed evolution stabilizes pBPARS, which likely underlies its improved performance.

To further test the scope of our optimized selections system, we sought to develop EcTyrRS mutants that charge additional useful ncAAs. One of the most popular classes of ncAAs are those with bioorthogonal conjugation handles such as an azide. EcTyrRS has been engineered to charge p-azidophenylalanine,^6, 10^ but this aromatic azide is susceptible to endogenous reduction. ^10, 33–34^ Engineering EcTyrRS to charge ncAAs with aliphatic azides will be beneficial, since these are significantly less prone to such undesirable loss. We pursued the selection of our EcTyrRS library using two different ncAAs, **2** and **3** (**Figure 1**), containing alkyl azides. Intriguingly, both selection yielded an overlapping set of mutants pAAFRS-6, 9, and 11 (**Figure 4A**), indicating that these mutants are likely polyspecific; i.e, able to charge structurally similar ncAAs. Indeed, further characterization revealed that pAAFRS-9 can charge charge both 2 and 3 selectively (**Figure 4B**).

**Figure 4:**
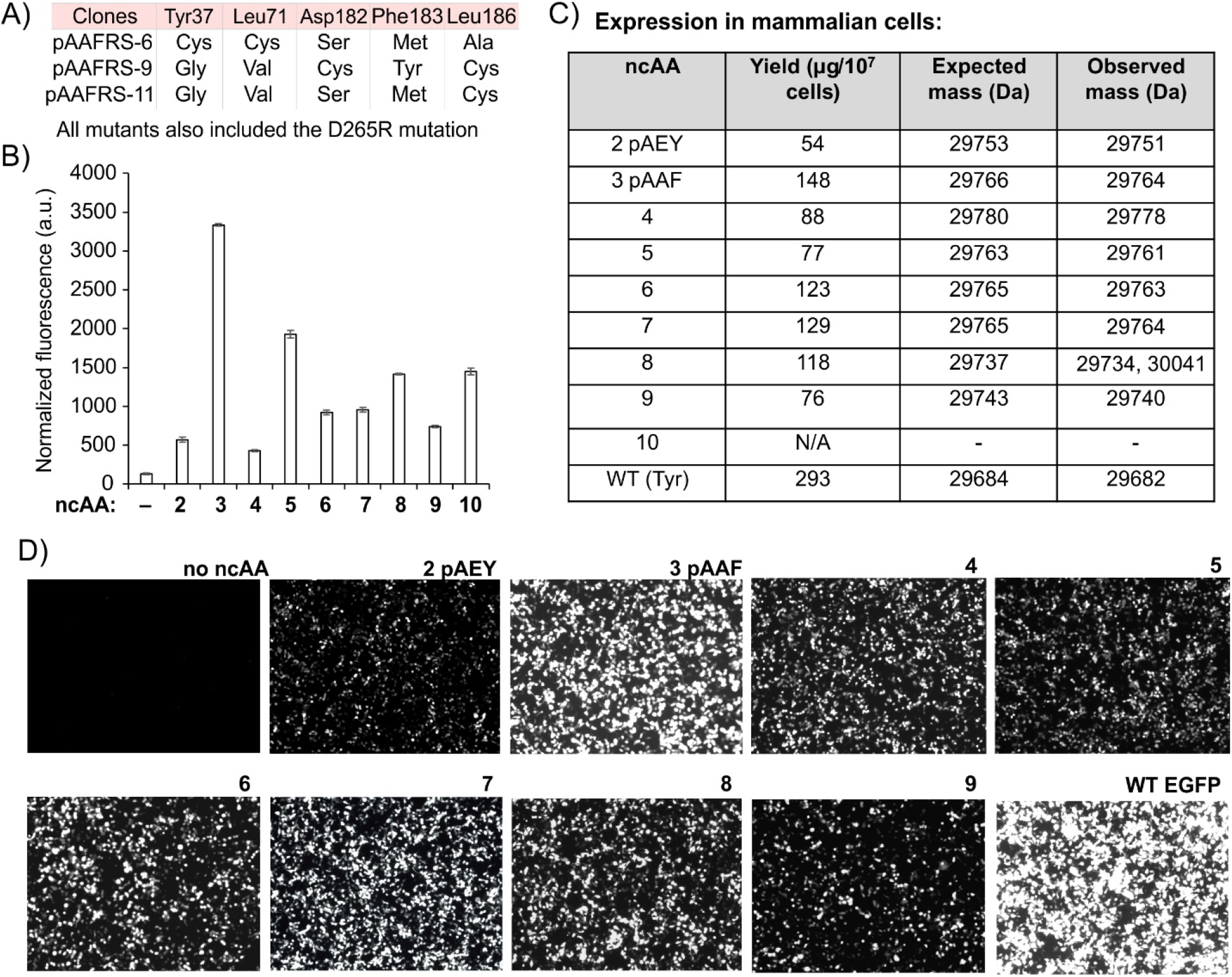
A highly polyspecific EcTyrRS mutant capable of charging ncAAs **2**-**10**. A) Mutations associated with EcTyrRS variants identified from the selection. B) Activity of pAAFRS-9 in ATMY E. coli strain, measured using sfGFP-151-TAG as the reporter. C) Incorporation of ncAAs 2-10 in HEK293T cells using EGFP-39-TAG as the reporter. Isolated yields for the reporter proteins and observed molecular weight of each are shown. D) Representative fluorescence images of HEK293T cells expressing the EGFP-39-TAG reporter shown in panel C.

Polyspecific aaRS mutants are valuable, as these enable rapid expansion of the ncAA toolbox with many new members, without having to perform individual selections for each.^10, 35^ To take advantage of the potential polyspecificity of pAAFRS-9, we synthesized several ncAAs that are structurally similar to its efficient substrate pAAF (Figure 1). These included additional bioconjugation handles such as alkyne and alkene (**4**-**7**) and electrophilic amino acids (**8**-**10**) which has been useful to forge proximity-dependent crosslink between interacting proteins,^36–37^ and for macrocyclization of genetically encoded peptides using a proximal cysteine residue.^38–39^ These substrates were found to be charged efficiently by pAAFRS-9 in both ATMY *E. coli* (**Figure 4B**) observed using a sfGFP-151-TAG reporter. We also tested the activity of pAAFRS-9 in mammalian cells using a EGFP-39-TAG reporter, and all but ncAA **10** were incorporated efficiently (**Figure 4C-D**). Although ncAA **10** could be used in ATMY *E. coli*, it showed high toxicity toward mammalian cells, preventing its incorporation therein. Incorporation of ncAAs **2-9** were confirmed by isolating the resulting EGFP-39-TAG reporter from HEK293T cell-free extract using immobilized metal-ion chromatography followed by mass-spectrometry. Of particular note is the observation that the aliphatic azide pAAF **3** shows no evidence of post-translational reduction (**Figure S5A**), providing a more robust bioconjugation handle relative to p–azidophenylalanine in eukaryotes. Also notable is the MS analysis of the reporter protein encoding the ncAA **8**, containing an electrophilic acrylamide group, which shows the presence of a major glutathione adduct (**Figure S5B**). Similar glutathione adducts have been observed before with the same ncAA incorporated into reporter proteins expressed in *E. coli* using an engineered archaeal TyrRS/tRNA pair.^36^ These enabling ncAAs are expected to significantly enhance the impact of the toolbox empowered by EcTyrRS engineering. Moreover, additional ncAAs with a similar structures would likely be accepted by pAAFRS-9, which we will continue to explore.

Tyrosine phosphorylation is an important post-translational modification in eukaryotes.^40–41^ In *E. coli*, non-hydrolyzable structural mimics of phosphotyrosine such as p-carboxymethylphenylalanine (pCMF **12**; **Figure 5A**) can be site-specifically incorporated using an engineered archaea-derived TyrRS/tRNA pair. Site-specific pCMF incorporation has been used in many studies to interrogate structural and functional consequences of phosphorylation of particular tyrosine residues, underscoring its utility.^42–48^ However, this ncAA cannot be currently incorporated into proteins expressed in eukaryotes (the archaeal Tyr pair is not orthogonal in eukaryotes). The ability to do so will be valuable, providing opportunities to interrogate consequences of tyrosine phosphorylation of eukaryotic proteins in their native environment. Genetically encoding polar amino acid (such as pCMF) has been challenging through engineering the pyrrolysyl pair, which is most widely used for ncAA incorporation in eukaryotes. In contrast, EcTyrRS represents a great starting point for developing a pCMF-specific aaRS, given the success of its archaeal counterpart with a similar active site.^42^ To explore this possibility, we subjected the aforementioned EcTyrRS library to our optimized selection system, hoping to identify a pCMF-specific mutant. Screening of surviving clones following the double-sieve selection led to the identification of a unique mutant (**Figure 5B**) that enables expression of the sfGFP-151-TAG reporter in ATMY *E. coli* only in the presence of pCMF (**Figure 5C**). Key mutations in pCMFRS include D182G and Y37H, which opens up room for the carboxymethyl group and likely interact with the negatively charged carboxylate, respectively. pCMFRS also facilitated efficient expression of the EGFP-39-TAG reporter in HEK293T cells, selectively in the presence of pCMF (**Figure 5D-E**). The reporter protein was isolated from mammalian cells by IMAC in good yield (45 μg/ 10^7^ cells) and characterized by SDS-PAGE and mass-spectrometry to confirm the incorporation of pCMF (**Figure 5F**). The ability to site-specifically incorporate this non-hydrolyzable mimic of phosphotyrosine into proteins expressed in eukaryotes, including mammalian cells will be a valuable tool to probe the consequences of tyrosine phosphorylation. Additionally, it further attests the utility of our optimized selection system for engineering EcTyrRS substrate specificity.

**Figure 5.**
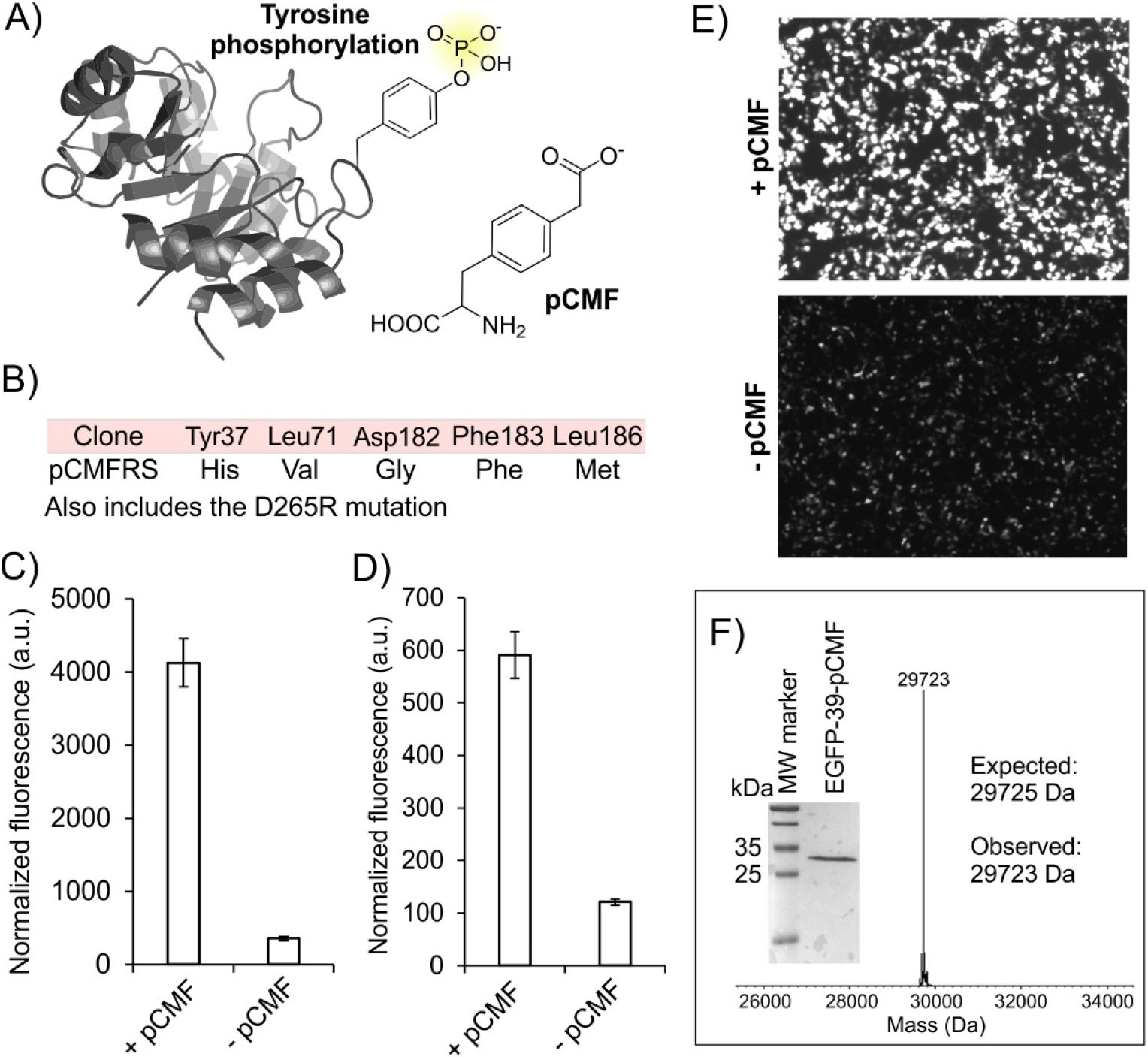
Genetically encoding pCMF in eukaryotes. A) pCMF mimics a phosphorylated tyrosine residue. B) Mutations associated with the pCMFRS identified through the selection of a EcTyrRS library. C) In ATMY E. coli, pCMFRS facilitates selective expression of the sfGFP-151-TAG reporter in the presence of pCMF. D) Activity of pCMFRS in HEK293T cells using the EGFP-39-TAG reporter demonstrate selective reporter expression in the presence of pCMF (fluorescence in cell-free extract). E) Representative fluorescence microscopy images of HEK293T cells expressing the EGFP-39-TAG reporter using pCMFRS in the presence and absence of pCMF. F) SDS-PAGE and ESI-MS analysis of the EGFP-39-pCMF reporter isolated from HEK293T cells.

In summary, we have developed an optimized *E. coli* based selection that enables facile engineering of EcTyrRS to introduce new chemistries into the genetic code of eukaryotes. Key to this selection system is an engineered tRNA_CUA_^EcTyr^ mutant, which we developed through directed evolution to exhibit little cross-reactivity in ATMY *E. coli*. We further developed a straightforward survival-based WiDR antibiotic selection that can differentiate between strongly and weakly active EcTyrRS mutants, and used it for significantly improving the activity of a first-generation poorly active pBPARS mutant through directed evolution. Together, our optimized selection system provides an opportunity to unlock the untapped potential of the EcTyrRS/tRNA^EcTyr^ pair for introducing previously inaccessible ncAAs into the genetic code of eukaryotes. We demonstrated the utility of our selection system by developing several new EcTyrRS mutants that allow efficient incorporation of several novel ncAAs that will be useful for several different applications. These include: 1) a highly efficient incorporation system for incorporating pBPA in mammalian cells, which is a popular photoaffinity probe, 2) several new ncAAs to facilitate bioconjugation, including pAAF, which is incorporated in mammalian cells with remarkable efficiency, and does not undergo reduction, 3) a series of electrophilic amino acid that can be used for proximal covalent cross-linking between interaction proteins, as well as for macrocyclization using a proximal cysteine residue, and finally, 4) a non-hydrolyzable analog of phosphotyrosine that will be useful for probing the consequences of this important post-translational modification. Finally, our work provides lessons that will be valuable for developing additional engineered aaRS/tRNA pairs for expanding the genetic code of eukaryotes using a similar strategy.

## ASSOCIATED CONTENT

### Supporting Information

Experimental methods, nucleotide sequences, supplementary figures and tables. (PDF)

## AUTHOR INFORMATION

### Notes

A patent application on the EcTyrRS mutants and ncAAs reported here has been submitted. AC is a cofounder and senior advisor of BrickBio, Inc., which focuses on applications of the ncAA mutagenesis technology

## ACKNOWLEDGMENT

This work was supported by NIH grant R35GM136437 to A.C.

