## Supporting information for "A facile platform to engineer *E. coli* tyrosyl-tRNA synthetase adds new chemistries to the eukaryotic genetic code, including a phosphotyrosine mimic"

**Supporting Figures**:

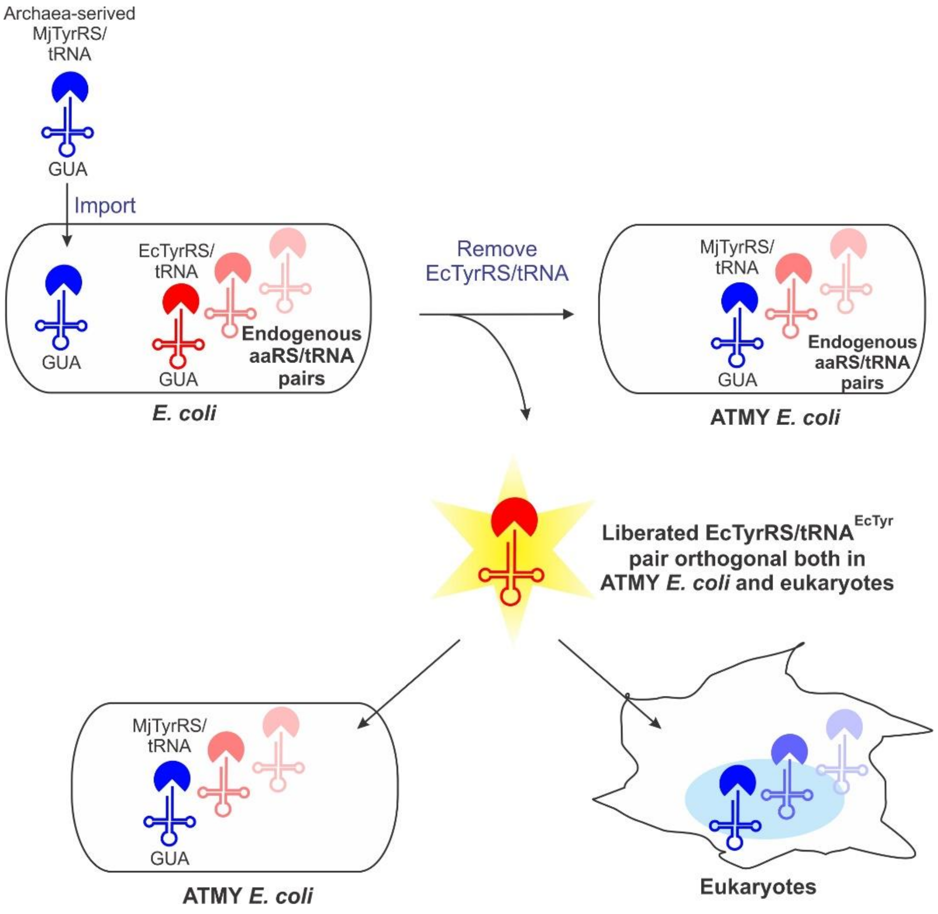


**Figure S1:** Scheme demonstrating the functional substitution of the endogenous EcTyrRS/tRNA pair in *E. coli* using an archaeal counterpart to generate ATMY strains. The liberated EcTyrRS/tRNA pair can be used as an orthogonal pair in both ATMY E. coli and eukaryotic cells. This enables engineering the substrate specificity of EcTyrRS using an *E. coli* based selection system in the ATMY strain, followed by the application of the resulting mutants in eukaryotic cells.

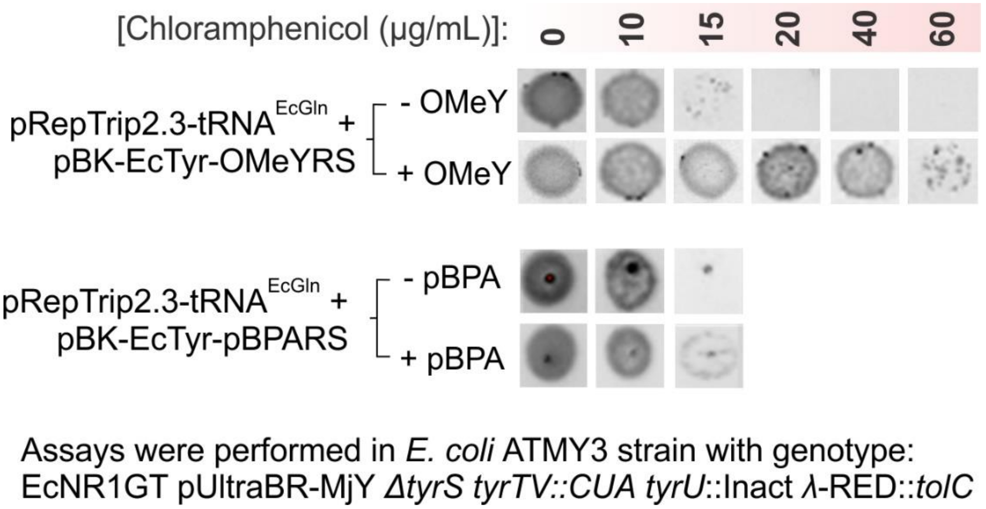


**Figure S2**: Background activity from the cross-charging of $tRNA_{CUA}^{EcTyr}$ by GlnRS masks the activity of weaker EcTyrRS mutants, such as pBPARS, preventing their selection. The assay is performed in ATMY3 strain, encoding a single copy of genomically integrated $tRNA_{CUA}^{EcTyr}$. The pRepTrip2.3-$tRNA^{EcGln}$ plasmid expresses a CAT reporter harboring a TAG at the permissive 98 site, as well as $tRNA^{EcGln}$, which attenuates the cross-reactivity between EcGlnRS and $tRNA_{CUA}^{EcTyr}$. In the absence of a cognate ncAA substrate for the EcTyrRS mutant, the cross-reactivity with GlnRS leads to survival at up to 10 µg/mL of chloramphenicol. In the presence of OMeYRS, which is a strongly active EcTyrRS mutant that charges OMeY, the cells survive up to 60 µg/mL of chloramphenicol. This activity is significantly higher than the background and is sufficient to enable its efficient selection.[1] However, weaker EcTyrRS mutants such as pBPARS, shows marginal activity over background (survival at up to 15 µg/mL), which is insufficient to facilitate their selection.

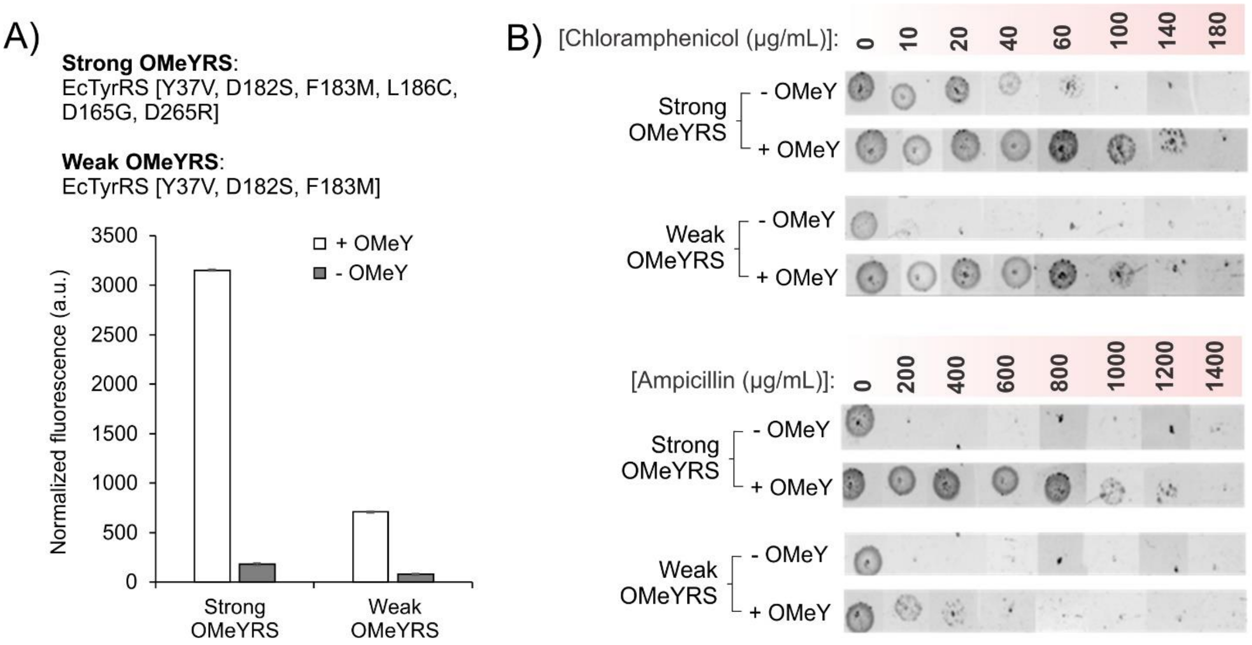


**Figure S3**: Development of a wide dynamic range (WiDR) antibiotic selection capable of differentiating between EcTyrRS mutants with high or moderate activity. A) Two established OMeYRS mutants used in this study, designated strong and weak. Using an sfGFP-151-TAG reporter in ATMY strain, we show that the strong OMeYRS facilitates approximately 6 fold higher reporter expression, relative to weak OMeYRS. B) Evaluating a chloramphenicol-acetyl transferase and a β-lactamase reporter, each encoding a TAG codon at a permissive site, for their ability to differentiate between the strong and weak OMeYRS. The CAT-98-TAG reporter, traditionally used for aaRS selection, shows a narrow dynamic range: the weak OMeYRS facilitates survival up to 100 μg/mL of chloramphenicol, while the strong variant supports up to 140 μg/mL. Such a narrow difference is unlikely to be suitable for serving as the basis of a successful selection of the strong variant. In contrast, use of the β-lactamase reporter resulted in the survival for the strong and the weak OMeYRS mutants at up to 1200 μg/mL and 400 μg/mL of ampicillin, respectively. The significantly wider dynamic range of the latter selection system makes it suitable for enriching more active EcTyrRS mutants.

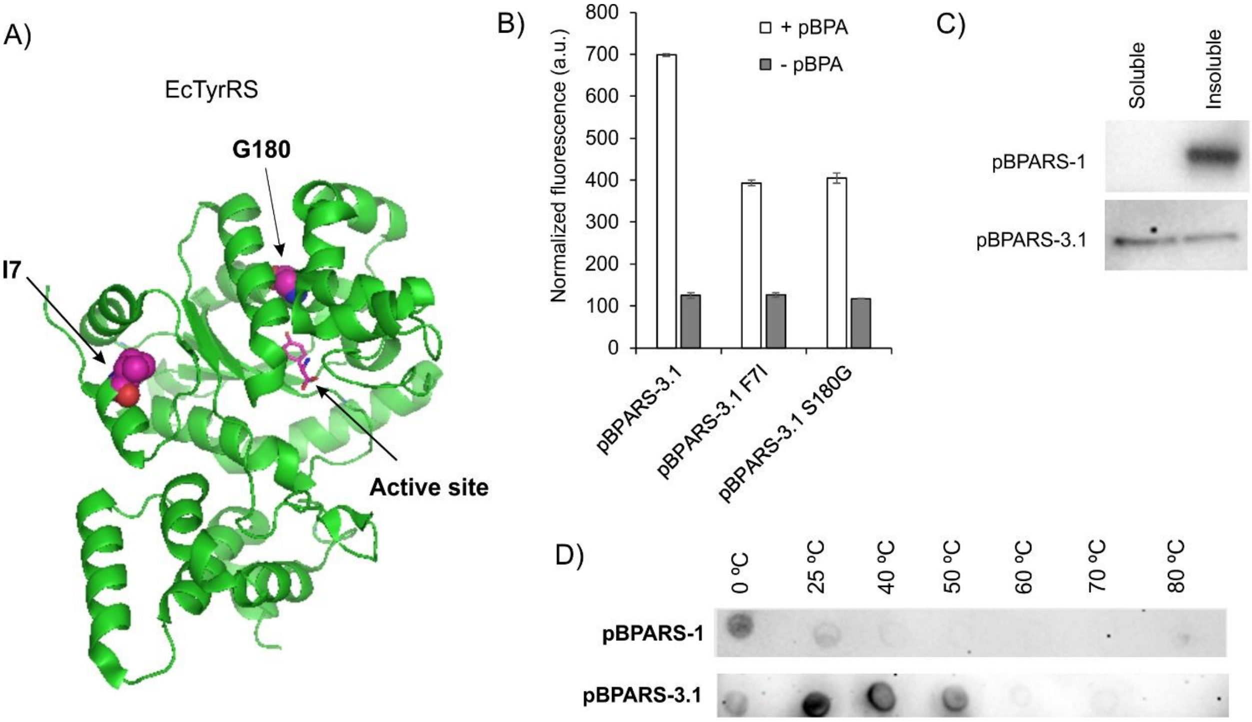


**Figure S4**: Mutations in pBPARS-3.1 enhances its thermostability. A) Crystal structure of EcTyrRS highlighting the sites of two key mutations: I7F and G180S. B) Reverting either I7F mutations attenuates the activity of pBPARS-3.1, suggesting both mutations contribute to its enhanced activity. Activities were measured in ATMY *E. coli* strain, using sfGFP-151-TAG as the reporter. C) When pBPARS-1 is expressed in ATMY E. coli, nearly all of it is found in the insoluble fraction of the cell-free extract, visualized by Western blot. In contrast, pBPARS3.1 is significantly more soluble. D) Thermostability assay in cell-free extract, where protein remaining in the soluble fraction after incubation at the target temperature is measured by immunoblotting, shows pBPARS-3.1 exhibits significantly higher robustness relative to pBPARS-1.

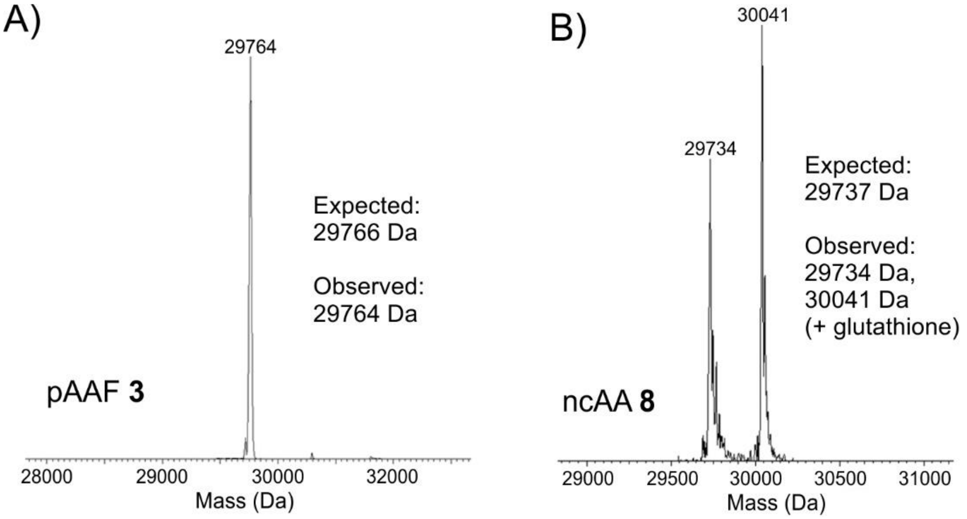


**Figure S5**: Deconvoluted ESI-MS analysis of EGFP-39-TAG reporter expressed in HEK293T cells using pAAFRS-9 in the presence of pAAF **3** (A) or ncAA **8** (B)

**Materials and methods**:

All cloning and plasmid propagation were performed in DH10B *E. coli* cells. Restriction enzymes, Phusion HS II High-Fidelity DNA polymerase, and IPTG were obtained from Fisher. T4 DNA ligase was obtained from Enzymatics. DNA extraction and PCR clean up were conducted with Macherey-Nagel Binding Buffer NTI and Epoch mini spin columns (Thermo-Fisher Scientific). Media components were obtained from Thermo-Fisher Scientific. DNA oligomers were purchased from Integrated DNA Technologies (IDT) and Genewiz was used for DNA sequencing. *E. coli* cultures were grown on LB agar plates or liquid medium with following antibiotic concentrations, unless otherwise noted: 90 μg/mL spectinomycin, 50 μg/mL chloramphenicol, 10 μg/mL gentamycin, 100 μg/mL ampicillin, 15 μg/mL zeocin, 12 μg/mL tetracycline, 30 μg/mL kanamycin. O-methyl-l-tyrosine was obtained from Fisher Scientific (AAH6309606), *p*-benzoyl-L -phenylalanine was purchased from Chem-Impex International (05110), pCMF was obtained from AsisChem (ASIS-0070). The rest of the ncAAs used here were chemically synthesized as described later.

**Safety Considerations**

No unexpected or new significant hazards were encountered.

**Strains, cell lines**. HEK293T cells (ATCC) were maintained at 37 $^{o}$C and 5% $CO_2$ in DMEM-high glucose (HyClone) supplemented with penicillin/streptomycin (Hyclone, final concentration of 100 U/L penicillin and 100 μg/mL streptomycin) and 10% fetal bovine serum (Corning).

The development of ATMY3, ATMY4, and ATMY5 *E. coli* strains are described in a previous publication by Italia *et al*.[1]

**Plasmids and cloning**:

pRepTrip2.3-EcOMeYRS was constructed by first PCR amplifying EcOMeYRS-VSML out of pBK EcOMeYRS-VSML[1] with the primers BKrep-SpeI-F and BKrep-BglII-R. The PCR product was digested with SpeI/BglII and inserted into the pRep vector backbone.

pRepTrip2.3p-EcYtR-h1 was created by PCR amplifying EcYtR-h1 from pBK EcYtR-h1 with BKrep-SpeI-F and BKrep-BglII-R. The PCR product was digested with SpeI/BglII and inserted into the pRep vector backbone.

pNeg-EcYtR-h1-barnase-2XTAG was created by PCR amplifying EcYtR-h1 from pBK EcYtR-h1 with NEGrep-SphI-F and BKrep-NcoI-R. The PCR product was digested with SphI/NcoI and inserted into the pNeg backbone that was digested with the same restriction enzymes.

All pB1U-EcTyrRS plasmids were generated by PCR amplifying the mutant EcTyrRS from its pBK plasmid with primers EcYRS-NheI-F and EcYRS-XhoI-R, followed by digestion with NheI/XhoI. The digested product was then inserted into the pB1U vector backbone using the same restriction sites.

The pBK-pBPARS-3.1 mutation reversion mutants were made through QuikChange mutagenesis (Agilent) with the following primers: pBPARS-F7I-F, pBPARS-S180G-F.

**Assessment of EcTyrRS-tRNA activity using a CAT-TAG reporter:**

5 mL LB media cultures of the ATMY strain (ATMY3 or ATMY5) harboring the appropriate pRepTrip2.3 (encoding $tRNA_{CUA}^{EcTyr}$ or $tRNA_{CUA}^{EcTyr}$-h1; also contains the CAT reporter with a Q98TAG mutation) and pBK-EcTyrRS were grown overnight from a single colony. Following overnight growth, the cultures were diluted to an $OD_{600}$ of 0.03, and 3 µl of this diluted culture was spotted on LB agar plates supplemented with kanamycin, spectinomycin, tetracycline, 0.02% arabinose, varying chloramphenicol concentrations, and +/- 1 mM of UAA. Growth on the plates was analyzed 24 and 48 hours following inoculation.

**Assessment of aaRS-tRNA activity using a sfGFP-151 reporter**

The pEvol-sfGFP-151-TAG reporter and the pBK-EcTyrRS plasmids were co-transformed into the ATMY4 (encoding two copies $tRNA_{CUA}^{EcTyr}$in the genome) for expression.[2] A 5 mL overnight culture was inoculated with a single colony from the transformation plates. Following overnight growth, 200 µL of the overnight culture was used to inoculate a 20 mL LB media culture supplemented with spectinomycin, kanamycin, and chloramphenicol. This culture was then grown at 37 $^{o}$C with shaking (250 rpm) to a final $OD_{600}$ of 0.6. At this point, the cultures were induced with IPTG (1 mM), the appropriate ncAA was added to the media, and incubated for 16 hours at 30 $^{o}$C with shaking (250 rpm). The cultures were then spun down (4,500 g, 10 min, 4 $^{o}$C), the media was removed, and the pelleted cells were resuspended in PBS. The resuspended cells were diluted 10 fold with PBS (15 µL resuspended cells added to 135 µL PBS) and fluorescence was measured in a 96 well-plate using a SpectraMAX M5 (Molecular Devices) (ex = 488 nm and em = 510 nm). Mean of three independent experiments were reported, and error bars represent standard deviation.

**Assessment of aaRS-tRNA activity using a EGFP-39-TAG reporter**

HEK293T cells were cultured in Dulbeco's modified Eagle's medium (high glucose DMEM) supplemented with 10% fetal bovine serum (FBS) and penicillin-streptomycin (0.5x) at 37 $^{o}$C in the presence of 5 % $CO_2$. The cells were seeded at a density of 600,000 cells per 12-well plate 24 hours prior to transfection. Co-transfection with pAcBac1-EGFP-39-TAG and pB1U aaRS was carried out when the cells were ~70% confluent. For the co-transfection, PEI (Sigma) and DNA were mixed at a ratio of 4 µL PEI (1 mg/mL) to 1 µg of total DNA (500 ng of each plasmid) in DMEM. This PEI/DNA mixture was incubated for 10 minutes at room temperature and then added to each well (100 µL per well). Then, ncAAs were added to each well (1 mM final concentration). Fluorescence images were taken 48 hours after transfection using a Zeiss Axio Observer fluorescence microscope.

**EGFP-39-TAG expression and purification**

The pAcBac1-EGFP-39-TAG and pB1U-EcTyrRS plasmids were co-transfected into HEK293T cells. One day before transfection, the HEK293T cells were seeded at a density of 8 x $10^6$ cells per 10 cm dish. Once the cells reached ~90% confluence, they were transfected using a transfection mixture of 50 µL PEI MAX (1 mg/mL; Polysciences) + 5 µg of each plasmid DNA

(10 µg total), and 200 µl DMEM. The transfection mixture incubated for 10 minutes prior to transfection and added to the 10 cm dish, followed by the ncAA (final concentration of 1 mM; 5 mM for pCMF). After 48 hours of expression, cells were harvested, washed twice with PBS buffer (spun at 5,000 g, 5 min, 4 °C), and lysed with CelLytic M lysis buffer (Sigma) supplemented with 1x Halt Protease Inhibitor and 0.01% Pierce Universal Nuclease. Following resuspension, the lysed cells were incubated at room temperature for 20 min. The cell-free extract was clarified then spun down (spun at 16,000 g, 5 min, 4 °C) and the EGFP reporter was purified using HisPur Ni-NTA resin following the manufacturer's protocol. Protein purity was characterized with both SDS-PAGE and whole protein ESI-MS ESI-MS (Agilent Technologies 6230 TOF LC/MS which is in-line with the 1260 Infinity HPLC system)

**Construction of the pBK $tRNA_{CUA}^{EcTyr}$ library**

Site-saturation mutagenesis was used to randomize five base-pairs of the acceptor stem of the $tRNA_{CUA}^{EcTyr}$ to create the library (theoretical diversity 1.05 x $10^6$). The $tRNA_{CUA}^{EcTyr}$ was amplified as two overlapping PCR fragments with Phusion HSII with the following primers: pBKseqtF+mutiR, mutiF+JI MCS sqR. The 5′ and 3′ fragments were joined together by primer-less overlap-extension PCR. The full-length PCR product was gel purified (1% agarose gel, 150 V) followed by amplification with the terminal primers pBKseqtF and JI MCS sqR. The PCR amplified insert was digested with BamHI/NcoI and ligated by T4 DNA Ligase into the pBK vector digested with the same restriction enzymes. The ligation mixture was ethanol precipitated with yeast-tRNA (Ambion) and transformed into DH10B electrocompetent cells. The library was covered using > $10^7$ distinct CFU.

**Selection for an orthogonal $tRNA_{CUA}^{EcTyr}$**

ATMY5 was co-transformed with the pBK $tRNA_{CUA}^{EcTyr}$ library and the positive selection reporter plasmid pRepTrip2.3-EcOMeYRS. This reporter construct contains a glnS-promoted *EcTyr*-OMeYRS, a CAT reporter mutagenized at Q98 to a TAG stop codon, an arabinose inducible T7 RNA polymerase mutagenized at positions 8 and 114 to stop codons (TAG), and T7-promoted wild-type GFPuv. Successful suppression of the TAG codons by active $tRNA^{EcTyr}{}_{CUA}$ library members leads to full-length expression of the CAT-reporter conferring chloramphenicol resistance. Additionally, suppression of the T7 RNA polymerase drives expression of GFPuv. 2.3 x $10^7$ CFU were plated on LB + Spec/Tet/Kan/Amp/0.02% arabinose + chloramphenicol (30 and 50 µg/mL) in the presence of 1 mM OMeY for 24 hours at 37 °C.

The pBK $tRNA_{CUA}^{EcTyr}$ plasmid from the surviving population was isolated and co-transformed into ATMY5 with the negative selection plasmid pNeg-noEcYtR. This reporter construct harbors a toxic barnase gene with stop codons at positions 1 and 90 under the control of an arabinose inducible promoter. 1.8 x $10^7$ CFU were plated on LB + Amp/Kan/Spec/0.2% arabinose in the absence of any ncAA for 12 hours at 37 °C.

Surviving pBK $tRNA_{CUA}^{EcTyr}$ plasmids from the negative selections were retransformed into ATMY5 with pRepTrip2.3-EcOMeYRS and plated under the positive selection condition. Individual clones from this plate were characterized for activity and cross-reactivity using the CAT-TAG reporter.

**pBPARS mutant library generation through error-prone PCR (ePCR)**

pBK-EcTyr-pBPARS-1 was used as the template to PCR amplify the pBPARS under error-prone conditions with the primers JI MCS sqR and pBKseqtF. TAQ-polymerase was used to amplify the insert under the following conditions: 0.3 or 0.15 M $MnCl_2$, 0.4 mM dCTP, 0.4 mM dTTP, 0.08 mM dATP, 0.08 mM dGTP, and 25 ng of DNA. The amplified pBPARS library was inserted into the pBK vector through PIPE cloning.[2] To execute the PIPE cloning, the pBK vector was pre-digested with NdeI/NcoI, agarose gel purified (1%), and amplified with the following primers: pBK-backbone-F and pBK-backbone-R. The vector and insert PCR products were then mixed at equal volumes and co-transformed into DH10B cells generating 4.5 x $10^6$ CFU.

**WiDR selection of ePCR-randomized pBPARS mutant library**

The pBK-pBPARS ePCR randomized library was transformed into ATMY5 containing the positive selection plasmid pRepTrip2.3-EcYtR-h1. This pRepTrip plasmid contains a proK-promoted mutant $tRNA_{CUA}^{EcTyr}$, a β-lactamase reporter mutagenized at the third amino acid to a stop codon (TAG), a CAT-Q98-TAG reporter, an arabinose inducible T7 RNA polymerase mutagenized at amino acids 8 and 114 to stop codons (TAG), and T7-promoted wild-type GFPuv. The incorporation of pBPA at the stop codons by active EcYRS-pBPARS ePCR library members leads to full-length expression of the β-lactamase conferring ampicillin resistance. Additionally, suppression of the T7 RNA polymerase drives expression of GFPuv. 5.7 x $10^7$ CFU were plated on LB + Spec/Tet/Kan /0.02% arabinose + 200 μg/mL ampicillin (400, 600, 800 and 1000 μg/mL) + 3 μg/mL chloramphenicol in the presence of 1 mM pBPA for 24 hours at 37 $^oC$. The pBK clones from the surviving cells were isolated, transformed into ATMY5 harboring pNeg-EcYtR-h1-barnase-2XTAG, and plated on LB + Spec/Amp/Kan/0.02% arabinose in the absence of pBPA. Finally, the surviving pBK plasmids were transformed into ATMY5 containing the positive selection plasmid pRepTrip2.3-$tRNA_{CUA}^{EcTyr}$-h1, and plated on LB-agar plates + Spec/Tet/Kan /0.02% arabinose + ampicillin (400, 600, 800 and 1000 μg/mL) + 3 μg/mL chloramphenicol in the presence of 1 mM pBPA. Clones surviving high ampicillin concentrations were characterized individually using the β-lactamase-3-TAG reporter, and sfGFP-151-TAG reporter.

**Construction of the pBK EcTyrRS library**

The EcYRS-lib1 was generated as described previously.[1] Overlap extension was used to introduce the D265R mutation into this library (theoretical diversity 1.06 x $10^7$).[2] Two PCR products of the EcYRS-lib1 were amplified with Phusion HSII with the following primers: BKrep-SpeI-F, EcYRS-D265R-R, EcYRS-D265R-F, and BkrepBglII-R. The two fragments were joined together by primer-less overlap extension followed by agarose gel purification. The purified product was amplified with the terminal primers BKrep-SpeI-F and BKrep-BglII-R and digested with NdeI/PstI. This digested product was ligated by T4 DNA ligase into a pBK vector digested with the same restriction enzymes. DNA from the ligation mixture was ethanol precipitated with yeast-tRNA (Ambion) and transformed into DH10B electrocompetent cells. The library was covered using > $10^8$ distinct CFU.

**Selection of ncAA-specific EcTyrRS mutants**

The pBK-*Ec*TyrRS library was subjected to a two-tier selection scheme of alternating positive and negative selection steps in ATMY5. The library was transformed into ATMY5 harboring pRepTrip2.3p-EcYtR-h1. The reporter plasmid contained one copy of *proK*-promoted *Ec*Tyr-tRNA$_{CUA}$-h1, a TAG-inactivated (at 98 position) CAT gene, a TAG-inactivated β-lactamase gene, an arabinose-inducible TAG-inactivated T7 polymerase, and a wild-type GFPuv reporter expressed from the *T7* promoter.[1] Suppression of the TAG codon would lead to chloramphenicol (Chlor) and ampicillin (Amp) resistance, as well as expression of the *T7*-promoted GFPuv. >$10^8$ colony forming units (cfu) were plated on LB agar containing spectinomycin (Spec), tetracyclin (Tet), and kanamycin (Kan), 0.02% arabinose, 30 µg/mL Chlor, 15 µg/mL carbenicillin, and 1 mM of the ncAA. Plates were incubated at 37 ˚C for 16-24 hours. Surviving colonies were harvested with LB and the surviving pBK plasmids were isolated.

The surviving pBK *Ec*TyrRS library members were co-transformed into electrocompetent ATMY5 cells harboring the negative selection reporter plasmid pNeg *Ec*YtRh1. The reporter plasmid contained an arabinose-induced barnase gene with two TAG codons (3TAG and 45TAG).[3] >$10^6$ cfu were plated on LB agar containing 1x Spec/Kan/Amp and 0.02% arabinose, and incubated at 37 °C for 12 hours. Surviving colonies were harvested and the pBK plasmids were isolated.

The pBK plasmids harvested from the negative selection were transformed back into electrocompetent ATMY5 cells containing pRepTrip2.3p-EcYtR-h1 for a second positive selection and plated on a chloramphenicol gradient containing 0, 15, 30, 45, 60 µg/mL Chlor in the presence and absence of 1 mM ncAA. All plates also contained Spec/Tet/Kan, 0.02% arabinose, and 15 µg/mL carbenicillin, and were incubated at 37 ˚C for 16 hours. Individual colonies screened for selective survival in the presence of the appropriate ncAA during the CAT-98-TAG assay. Individual colonies were inoculated into 0.5 mL LB with 1x Spec/Tet/Kan in a deep-well polypropylene plate and incubated overnight at 37 ˚C, shaking at 250 RPM. The overnight cultures were diluted to an $OD_{600}$ of 0.03 and 3 µL of each was spot plated on LB agar plates with the positive selection conditions in the presence or absence of the ncAA. Clones with the most prominent UAA-dependent growth were picked, sequenced, and spot-plated again to confirm the growth phenotype.

**Table S1**: Oligonucleotide sequences.

| Primer name | Sequence |
|---|---|
| **BKrep-BglII-R** | AATAATAagatctGCTCCTTAGATCTTCCTAGGACCATTCC |
| **BKrep-NcoI-R** | AATAATAccatggGCTCCTTAGATCTTCCTAGGACCATTCC |
| **BKrep-SpeI-F** | AATAATAactagtATTACGCTGACTTGACGGGACGG |
| **BKrep-BamHI-F** | AATAATAggatccGCGCTTGTTTCGGCGTGGGTATG |
| **EcYRS-D265R-F** | GATCAACACTGCGCGTGCCGACGTTTACCGCTTCCTGAAGTTCTTCAC |
| **EcYRS-D265R-R** | GTGAAGAACTTCAGGAAGCGGTAAACGTCGGCACGCGCAGTGTTGATC |
| **EcYRS-NheI-F** | TTTGAGGAATCCGCTAGCGCAAGCAGTAACTTGATTAAACAATTGCAAGAG |
| **EcYRS-XhoI-R** | AATTCTCGAGTTATTTCCAGCAAATCAGACACTAATTC |
| **JI MCS sqR** | GAGATCATGTAGGCCTGATAAGCGTAGC |
| **mutiF** | cAAAGGGAGCAGACTCTAAATCTGCCGTCATCGACTTCGAAGGTTCGAATCCTTC C**NNNNN**CACCACTGCAGATCCTTAGCGAAAGCTAAG |
| **mutiR** | cTTCGAAGTCGATGACGGCAGATTTAGAGTCTGCTCCCTTTGGCCGCTCGGGAAC **NNNNN**CGAATTCAGCGTTACAAGTATTACACAAAGTTTTTTATG |
| **NEGrep-SphI-F** | AATAATAgcatg**c**TCGAACTTTTGCTGAGTTGAAGG |
| **pBKseqtF** | AATAATA**gcatgc**TCGAACTTTTGCTGAGTTGAAGG |
| **pBPARS-F7I-F** | ATGGCAAGCAGTAACTTGATTAAACAATTGCAAGAGCGGG |
| **pBPARS-S180G-F** | GTTTTCCTACAACCTGTTGCAGGGTTATGGGTTC |

### Synthesis of the non-canonical amino acids (ncAA):

### General information:

All reactions were carried out with anhydrous solvents (obtained from Acros Organics, Fisher Scientific or Alfa Aesar) under an atmosphere of $N_2$ in oven dried glasswares. Boc-4-Amino-*L*-phenylalanine was obtained from Combi-Blocks; 4-Amino-L-phenylalanine was obtained from Santa Cruz Biotechnology. All other reagents were obtained from Fischer Scientific or Sigma-Aldrich and used without further purification. All work up and purifications were carried out with reagent grade solvents under ambient conditions. Room temperature refers to 22 °C. NMR spectra were recorded on Varian INOVA 500 MHz or 600 MHz NMR spectrometers at 25 °C. NMR data were processed using Mnova (Mestrelab). Chemical shifts are reported in ppm from tetramethylsilane (δ 0.00) with the solvent resonance as the internal standard. For $^{1}$H-NMR, data are reported as follows: chemical shift, multiplicity (s = singlet, d = doublet, dd = double doublet, t = triplet, m = multiplet), coupling constants (Hz) and integration. All the reactions were monitored by liquid chromatography (Agilent Technologies 1260 Infinity) coupled mass-spectrometry (Agilent Technologies 6230 TOF LC/MS) using a reverse phase analytical column with mobile phase comprises of water-acetonitrile-0.1% formic acid.

The non-canonical amino acid stock solutions (30 mM-100 mM) were adjusted to pH 7-8 and sterile filtered through a 0.22 µm syringe filter (source: Fisherbrand) before adding to the corresponding cell culture.

### Synthesis of ncAA 2:

The ncAA **2** was prepared following a literature procedure[3] through the following synthetic sequence:

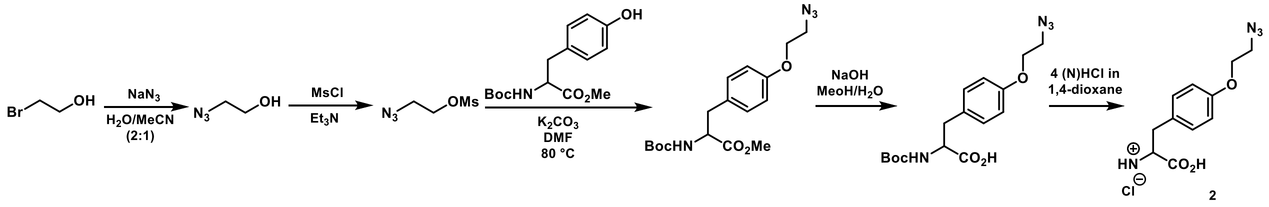


### Synthesis of 3 (pAAF):

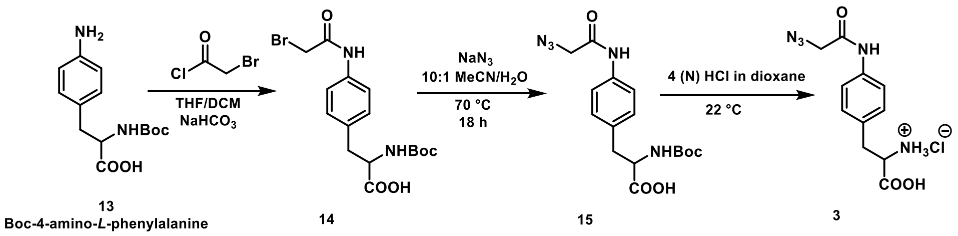

To a solution of Boc-4-Amino-*L*-phenylalanine (2.4 g, 8.56 mmol) in 50 mL THF/$CH_2Cl_2$ (4:1) mixture sodium bicarbonate (4.1 g, 48.8 mmol) was added. Resulting heterogeneous mixture was cooled in an ice-salt bath and bromoacetyl chloride (3.5 mL, 42 mmol) was added dropwise over 5 min while vigorously stirring under an inert atmosphere. After overnight stirring at the room temperature full consumption of Boc-4-Amino-*L*-phenylalanine was observed by LC-MS analysis of the crude reaction mixture. The mixture was filtered through a fritted funnel using 200 mL EtOAc. The filtrate was chilled on ice and washed sequentially with 100 mL chilled aqueous 0.5 (N) HCl (*caution*: vigorous effervescence), water (30 mL) and brine (30 mL). The organic layer was separated. The combined aqueous layer (pH 5-6) was re-extracted with ethyl acetate (2x75 mL); combined organic layer was washed with water (30 mL) and brine (30 mL). The organic layers were pooled, dried over anhydrous $Na_2SO_4$ and concentrated under reduced pressure to get the N-Boc protected α-bromoamide intermediate (**14**) as an oil (3.1 g, 90% yield; >95% pure by LC-MS) that eventually solidified at −20 °C. The intermediate was used directly in the next step without further purification.

The crude solid of α-bromoamide (**14**; 2 g, 4.98 mmol) was dissolved in 30 mL MeCN and 4.5 g $NaN_3$ was added followed by the addition of 3 mL water. The resulting heterogeneous mixture was stirred for 18 h at 70 °C under an inert atmosphere. Analysis of the crude reaction mixture by LC-MS showed full conversion. The mixture was allowed to come to the room temperature and concentrated under reduced pressure. The resulting semi-solid mass was diluted in a biphasic mixture of 200 mL EtOAc and 50 mL water; resulting aqueous layer should have pH 5-6. The Aqueous layer was washed off and organic layer was subsequently washed with water (2x40 mL) followed by brine (40 mL). The organic layer was dried over anhydrous $Na_2SO_4$ and concentrated under reduced pressure to get the Boc-protected α-azidoamide intermediate as an oil (1.8 g, quantitative yield, >95% pure by LC-MS) which was used in the next step without further purification.

Anhydrous 4 (N) HCl solution in 1,4-dioxane (20 mL) was added to the oil under an inert atmosphere. Resulting heterogeneous mixture was stirred for 4 h at the room temperature. LC-MS analysis of the crude mixture confirmed completion of the removal of the Boc protection. The solvent was evaporated off under reduced pressure to get an oil that was repeatedly diluted with 15 mL chloroform and dried under reduced pressure to get rid of dissolved hydrochloric acid. The resulting oil was dissolved in 5 mL of MeOH and then diluted with 230 mL diethylether. The turbid solution was allowed to settle for 15 h in the freezer (−20 °C). The solid precipitate was collected by filtration and dried under reduced pressure to get the hydrochloride salt of **3** (1.5 g, quantitative yield) as a white solid. **$^1$H-NMR (500 MHz, $D_2O$):** 7.45 (d, *J* = 10 Hz; 2H), 7.33 (d, *J* = 10 Hz; 2H), 4.30 (t, *J* = 5 Hz; 1H), 4.17 (s; 2H), 3.34 (dd, *J* = 5, 15 Hz; 1H), 3.21 (dd, *J* = 5, 15 Hz; 1H); **$^{13}$C-NMR (125 MHz, $D_2O$):** δ 171.65, 169.10, 135.77, 131.56, 130.11, 122.65, 54.25, 52.16, 35.14. **ESI-TOF-MS** (positive): m/z $[M+H]^+$ Calculated for $C_{11}H_{14}N_5O_3$ 264.1096 found 264.1047.

### Synthesis of ncAA 4:

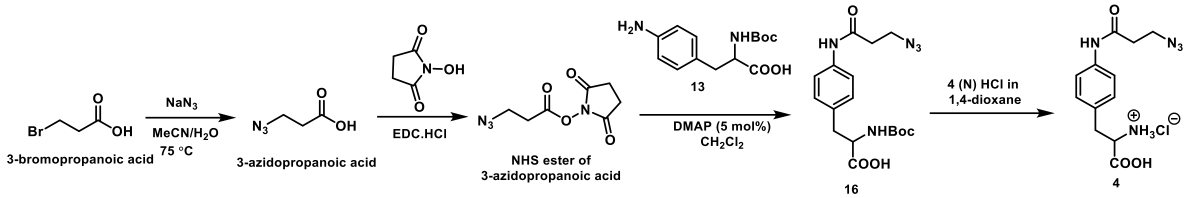


A mixture of 3-bromopropionic acid (4 g, 26.3 mmol) and 6 g $NaN_3$ was heated in 50 mL MeCN/water (4:1) mixture for 16 h at 75 °C. The mixture was allowed to come to the room temperature, concentrated under reduced pressure, acidified to pH 2 with aqueous 1 (N) HCl and extracted with EtOAc (3x75 mL). The organic layer was dried over anhydrous $Na_2SO_4$ and concentrated under reduced pressure to get 3-azidopropionic acid (quantitative yield) as an amber color oil which was used in the next step without further purification.

The oil (1.5 g, 13 mmol) was dissolved in 75 mL $CH_2Cl_2$ and N-hydroxysuccinimide (1.5 g, 13 mmol) was added followed by the addition of EDC.HCl (2.74 g, 14.25 mmol). The resulting mixture was stirred for 18 h at the room temperature and then washed sequentially with water (4X25 mL) and brine (30 mL). The organic layer was dried over anhydrous $Na_2SO_4$ and concentrated under reduced pressure to get the N-hydroxysuccinimide (NHS) ester of 3-azidopropionic acid (2.6 g, 94% yield) as an oil that eventually solidified in the freezer (−20 °C). The NHS ester was used in the next step without further purification.

The NHS ester of 3-azidopropionic acid (681 mg, 3.2 mmol) and 4-dimethylaminopyridine (DMAP, 23 mg, 5 mol%,) were sequentially added to a solution of N-Boc-4-Amino-*L*-phenylalanine (1g, 3.56 mmol) in 30 mL $CH_2Cl_2$. The resulting mixture was stirred at the room temperature for 24 h. On full conversion of the Boc-4-Amino-*L*-phenylalanine (judged by LC-MS), the mixture was diluted with 40 mL $CH_2Cl_2$. And the organic layer was washed sequentially with pre-chilled 0.1 (N) aqueous HCl solution (2x25 mL), 30 mL water and 30 mL brine. The combined organic layer was dried over anhydrous $Na_2SO_4$ and concentrated under reduced pressure to get the Boc-protected β-azidoamide intermediate **16** as an oil (1.10 g, 92% yield, >95% pure by LC-MS) that was used in the next step without further purification.

To the crude oil, an anhydrous 4 (N) HCl solution in 1,4-dioxane (15 mL) was added under an inert atmosphere and the resulting heterogeneous mixture was stirred at the room temperature for 5 h to get full Boc-deprotection (determined by LC-MS). The solvent was evaporated off in a under reduced pressure. The resulting oil was repeatedly diluted with 15 mL chloroform and evaporated to get rid of dissolved hydrochloric acid. The residue was dissolved in 5 mL of MeOH and then diluted with 230 mL diethylether. Resulting turbid solution was allowed to settle for 15 h inside a freezer (−20 °C) to get a white precipitate that was collected by decanting off the supernatant; drying off the remaining supernatant (mother liquor) under reduced pressure provided the hydrochloride salt of **4** (1g, quantitative yield) as a light brown solid. **$^1$H-NMR (500 MHz, $D_2O$):** 7.46 (d, $J$ = 10 Hz; 2H), 7.33 (d, $J$ = 10 Hz; 2H), 4.29 (dd, $J$ = 5, 5.5 Hz; 1H), 3.68 (t, $J$ = 5 Hz; 2H), 3.34 (dd, $J$ = 5, 15 Hz; 1H), 3.21 (dd, $J$ = 10, 15 Hz; 1H), 2.72 (t, $J$ = 5 Hz; 1H); **$^{13}$C-NMR**

**(125 MHz, $D_2O$):** δ 172.39, 171.71, 136.28, 131.24, 130.08, 122.41, 54.29, 47.15, 35.65, 35.15. **ESI-TOF-MS** (positive): m/z $[M+H]^+$ Calculated for $C_{12}H_{16}N_5O_3$ 278.1253 found 278.1177.

### Synthesis of ncAA 5:

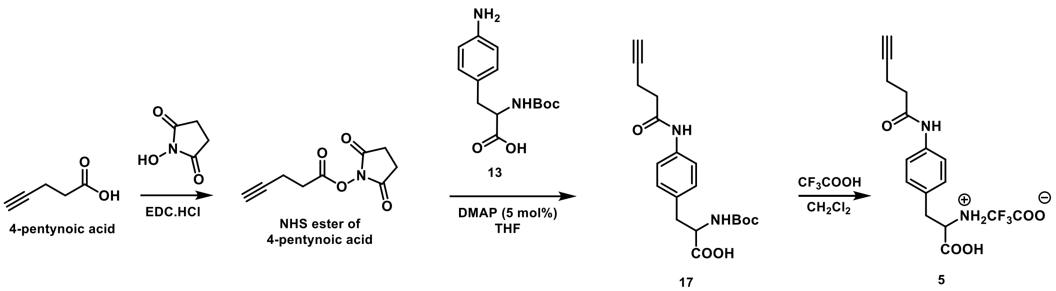


To a solution of 4-pentynoic acid (1 g, 10.2 mmol) in 50 mL $CH_2Cl_2$ N-hydroxysuccinimide (1.34 g, 11.6 mmol) was added followed by EDC.HCl (2.3 g, 12 mmol). The resulting mixture was stirred for 20 h at the room temperature under an inert atmosphere and then washed sequentially with water (4X20 mL) and 20 mL brine. The organic layer was dried over anhydrous $Na_2SO_4$ and concentrated under reduced pressure to get the N-hydroxysuccinimide (NHS) ester of 4-pentynoic acid as a colorless oil (1.8 g, 90% yield) that was used directly in the next step without further purification.

The N-hydroxysuccinimide ester of 4-pentynoic acid (526 mg, 2.69 mmol) and 4-dimethylaminopyridine (16 mg, 5 mol%) were sequentially added to a suspension of Boc-4-amino-*L*-phenylalanine (808 mg, 2.88 mmol) in 30 mL THF. The resulting heterogeneous mixture was stirred at 50 °C for 15 h to get the full conversion of the Boc-4-Amino-*L*-phenylalanine (judged by LC-MS). The solvent was evaporated off and the residue was dissolved in 75 mL $CH_2Cl_2$. The organic layer was washed sequentially with chilled 0.1 (N) aqueous HCl solution (2x25 mL), water (30 mL) and brine (30 mL). The combined organic layer was dried over anhydrous $Na_2SO_4$ and concentrated under reduced pressure to get the Boc-protected alkyne-amide intermediate **17** as an oil (852 mg, 88% yield, >95% pure by LC-MS) that was used in the next step without further purification.

The oil was dissolved in 10 mL dichloromethane and. chilled on an ice-salt bath (0 °C) under an inert atmosphere. To the resulting well stirred mixture 10 mL trifluoroacetic acid (anhydrous, HPLC grade) was added dropwise. The resulting mixture was stirred overnight at 4 °C and then concentrated under reduced pressure at 30 °C. The oily residue was repeatedly diluted with 15 mL $CHCl_3$ and evaporated to get rid of trace trifluoroacetic acid. The resulting oil was dissolved in 5 mL MeOH and then diluted with 230 mL diethylether. The turbid solution was allowed to settle for 15 h in the freezer (−20 °C). The precipitation was collected by filtration and dried under reduced pressure to get the trifluoroacetate salt of **5** as a brown solid (880 mg, quantitative yield).**$^1$H-NMR (600 MHz, $D_2O$):** 7.48 (d, $J$ = 12 Hz; 2H), 7.35 (d, $J$ = 12 Hz; 2H), 4.29-4.26 (m; 1H), 3.38 (s; 1H), 3.37-3.33 (m; 1H), 3.23-3.20 (m; 1H), 2.68-2.65 (m; 2H), 2.61-2.59 (m; 2H); **$^{13}$C-NMR (150 MHz, $D_2O$):** δ 176.14, 174.76, 138.97, 134.00, 132.75, 125.08, 85.63, 72.84,

57.26, 37.94, 37.69, 17.05. **ESI-TOF-MS** (positive): m/z $[M+H]^+$ Calculated for $C_{14}H_{17}N_2O_3$ 261.1239 found 261.1095.

### Synthesis of ncAA 6:

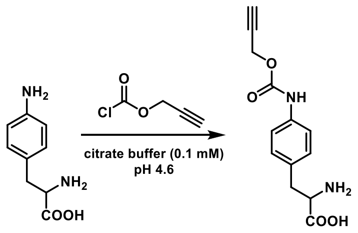


6

In a vigorously stirred solution of 4-Amino-*L*-phenylalanine (500 mg, 2.8 mmol) in 10 mL citrate buffer (100 mM, pH 4.6), propargyl chloroformate (275 μL; 2.8 mmol). was added dropwise. As a solid started precipitating out, the pH was adjusted to pH 4-4.6 with dropwise addition of aqueous 2 (N) NaOH during the progress of the reaction. After 3 h of stirring at the room temperature, the heterogeneous mixture was diluted with 30 mL of ice cold citrate buffer and the solid precipitate was collected by centrifugation (5000 rcf, 10 min) in a 50 mL conical tube at 4 °C. The solid residue was washed with 10 mL cold water and dried thoroughly to get the product **6** as a beige color solid. **$^1$H-NMR (500 MHz, DMSO-D6):** 9.78 (s; 1H), 7.34 (d, *J* = 10 Hz; 2H), 7.17 (d, *J* = 10 Hz; 2H), 4.72 (s; 2H), 3.41-3.38 (m; 2H), 3.09-3.05 (m; 1H), 2.83-2.78 (m; 1H); **$^{13}$C-NMR (125 MHz, DMSO-D6):** δ 170.18, 153.11, 137.64, 132.08, 130.19, 118.87, 79.46, 77.91, 56.01, 52.30, 36.57. **ESI-TOF-MS** (positive): m/z $[M+H]^+$ Calculated for $C_{13}H_{15}N_2O_4$ 263.1031 found 263.0970.

### Synthesis of ncAA 7:

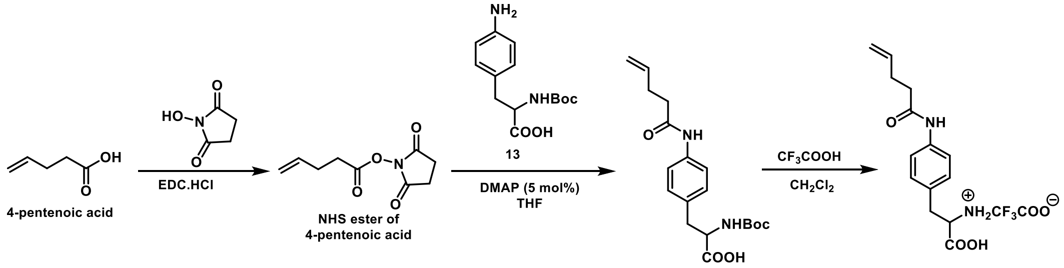


18 7

N-hydroxysuccinimide (2.6 g, 22.6 mmol) was added to a solution of 4-pentenoic acid (2 g, 20 mmol) in 70 mL $CH_2Cl_2$ followed by the addition of EDC.HCl (4.6 g, 24 mmol). The resulting mixture was stirred for 20 h at the room temperature under an inert atmosphere and then washed sequentially with water (4X25 mL) and 25 mL brine. The organic layer was dried over anhydrous $Na_2SO_4$ and concentrated under reduced pressure to get the N-hydroxysuccinimide (NHS) ester of 4-Pentenoic acid as a colorless oil (3.58g, 91% yield).

The N-hydroxysuccinimide ester (600 mg, 3 mmol) and 4-dimethylaminopyridine (16 mg, 5 mol%) were sequentially added to a suspension of N-Boc-4-amino-*L*-phenylalanine (900 mg, 3.21 mmol) in 30 mL dichloromethane. The resulting heterogeneous mixture was stirred at the room temperature for 24 h to get the full conversion (judged by LC-MS). The mixture was diluted with 40 mL $CH_2Cl_2$ and washed sequentially with chilled 0.1 (N) aqueous HCl solution (2x25 mL), water (30 mL) and brine (30 mL). Combined organic layer was dried over anhydrous $Na_2SO_4$ and concentrated under reduced pressure to get the Boc-protected alkene-amide intermediate **18** as an oil (934 mg, 86% yield, >95% pure by LC-MS) that was used in the next step without further pufication.

The oil was dissolved in 10 mL dichloromethane and. chilled on an ice-salt bath (0 °C) under an inert atmosphere. To the resulting well stirred mixture 10 mL trifluoroacetic acid (anhydrous, HPLC grade) was added dropwise. The resulting mixture was stirred at 4 °C overnight and then concentrated under reduced pressure at 30 °C. The oily residue was repeatedly diluted with 15 mL $CHCl_3$ and evaporated to get rid of trace trifluoroacetic acid. The resulting oil was dissolved in 5 mL MeOH and then diluted with 230 mL diethylether. Resulting turbid solution was allowed to settle for 15 h in the freezer (−20 °C). The solid precipitation was collected by suction filtration to get the desired product **7** as a brown solid (970 mg, quantitative yield). **$^1$H-NMR (500 MHz, DMSO-D6):** 9.99 (s; 1H), 7.54 (d, *J* = 10 Hz; 2H), 7.17 (d, *J* = 10 Hz; 2H), 5.88-5.80 (m; 1H), 5.08-5.04 (m; 1H), 4.98-4.96 (m; 1H), 4.01 (t, *J* = 5; 1H), 3.09-2.99 (m; 1H), 2.41-2.38 (m; 2H), 2.35-2.30 (m; 2H); **$^{13}$C-NMR (125 MHz, DMSO-D6):** δ 170.95, 170.93, 138.79,137.98, 130.16, 130.06, 119.63, 115.61, 54.13, 35.90, 29.52. **ESI-TOF-MS** (positive): m/z $[M+H]^+$ Calculated for $C_{14}H_{19}N_2O_3$ 263.1395 found 263.1434.

**<u>Synthesis of ncAA 8:</u>**

ncAA **8** was prepared following a literature procedure[4] through the following synthetic sequence:

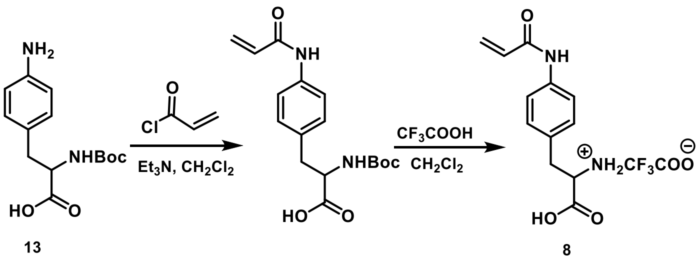

### Synthesis of ncAA 9:

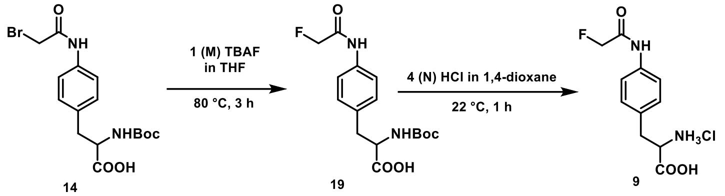


To the N-Boc protected α-bromoamide intermediate (**14**; 500 mg, 1.24 mmol; preparation described above) a solution of anhydrous tetrabutylammonium fluoride (TBAF) solution (1 M) in tetrahydrofuran (2.5 mL, 2.5 mmol) was added dropwise. The mixture was stirred at 80°C for 2 h and then allowed to come to the room temperature. LC-MS showed complete conversion of the α-bromoamide intermediate to the N-Boc protected α-fluoroamide intermediate. The resulting mixture was diluted with ethyl acetate (50 mL) and then washed with water (3x20 mL). The organic layer was washed with brine (20 mL), dried over anhydrous sodium sulfate and concentrated under reduced pressure to afford the desired product N-Boc protected α-fluoroamide as an oil (340 mg, 81% yield) that was used in the next step without further purification.

An anhydrous 4 (N) HCl solution in 1,4-dioxane (5 mL) was added dropwise to the oil (N-Boc α-fluoroamide intermediate) under an inert atmosphere. The resulting heterogeneous mixture was stirred at the room temperature for 1 h to have the complete deprotection of the Boc group. Dry ether (10 mL) was added to the reaction mixture and the supernatant was decanted off. The residue was repeatedly titurated with chloroform and then thoroughly dried to afford the salt of ### as a yellow color solid (210 mg, 75% yield). **$^{1}$H-NMR (600 MHz, $D_2O$):** 7.49 (d, $J$ = 12 Hz; 2H), 7.35 (d, $J$ = 12 Hz; 2H), 5.02 (d, $J_{H-F}$ = 48 Hz; 2H), 4.34-4.32 (m; 1H), 3.36-3.33 (m; 1H), 3.24-3.21 (m; 1H); **$^{13}$C-NMR (150 MHz, $D_2O$):** δ 174.12, 171.86 (d, $J_{C-F}$ = 18 Hz), 137.93, 134.30, 132.78, 125.49, 82.5 (d, $J_{C-F}$ = 180 Hz), 56.76, 37.74. **ESI-TOF-MS** (positive): m/z $[M+H]^+$ Calculated for $C_{11}H_{14}FN_2O_3$ 241.0988 found 241.0886.

### Synthesis of ncAA 10:

ncAA **10** was prepared following a literature procedure[5] through the following synthetic sequence:

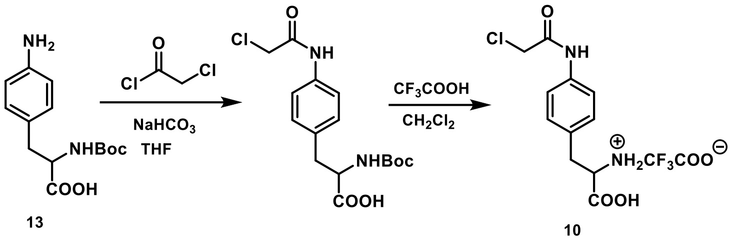


13

10
